# Survival of *Campylobacter jejuni* in *Acanthamoebae castellanii* provides mechanistic insight into host pathogen interactions

**DOI:** 10.1101/2022.07.26.501518

**Authors:** Fauzy Nasher, Burhan Lehri, Megan F Horney, Richard A Stabler, Brendan W Wren

**Author notes:** Address correspondence to **Brendan W. Wren**, and **Fauzy Nasher**.

## Abstract

*Campylobacter jejuni* is the leading cause of bacterial foodborne gastroenteritis world-wide but is rarely transferred between human hosts. Although a recognized microaerophile, *C. jejuni* is incapable of growing in an aerobic environment. The persistence and transmission of this pathogen outside its warm-blooded avian and mammalian hosts is poorly understood. *Acanthamoebae species*, are predatory protists and form an important ecological niche with several bacterial species. Here, we investigate the interaction of *C. jejuni* and *Acanthamoebae castellanii* at the single-cell level. We observe that a subpopulation of *C. jejuni* cells can resist killing by *A. castellanii* and non-digested bacteria are released into the environment where they can persist. In addition, we observe that *A. castellanii* can harbor *C. jejuni* even upon encystment. Transcriptome analyses of *C. jejuni* interactions revealed similar survival mechanisms when infecting both *A. castellanii* and warm-blooded hosts. In particular, nitrosative stress defense mechanisms and flagellum function are important as confirmed by mutational analyses. This study describes a new host-pathogen interaction for *C. jejuni* and confirms that amoebae are transient hosts for the persistence, adaptability and potential transmission of *C. jejuni*.

## Introduction

*Campylobacter jejuni* is the most prevalent gastrointestinal bacterial pathogen worldwide but is rarely transferred between human hosts suggesting that its high frequency in infection must relate to survival outside its warm-blooded hosts [1]. It is still ambiguous how this bacterium, an obligate microaerophile, remains omnipresent in the environment, given its inability to grow under atmospheric conditions [1, 2]. Although not all ‘environments’ out-side of the host may necessarily be aerobic, *C. jejuni* has been shown to form biofilms in atmospheric conditions as a mode of persistence [3, 4] and has also been recognized to transform into viable but non-culturable (VBNC) state [5, 6]. This allows the bacterium to withstand many stresses and remain viable with low metabolic activity for months [7]. Additionally, several studies have investigated *C. jejuni* environmental survival out-side of the host with free-living protists [8–12], however, the mechanistic understanding of this important niche remains poorly understood.

Free-living bacteria populations face constant challenges, particularly relating to predation by protists such as *Acanthamoebae* species [13–15]. Some bacteria have developed mechanisms to defend against killing within amoebic cells [14]. *Acanthamoebae spp*. have also been shown to have an important role in harboring and dissemination of pathogenic bacteria [13, 14, 16]. These interactions have been linked to several disease outbreaks of contaminated water [16] and food sources, including livestock [17]. *Acanthamoebae spp*. are characterized by a biphasic life cycle; a ‘trophozoite’ metabolically active form, and a ‘cyst’ state, which is a non-dividing dormant form [18]. Encystment occur from the trophozoite form, with changes to the cell morphology including; cell rounding and thicker cell-wall [19]. The thickened cell wall enhances resistance to disinfection treatments and environmental stresses, which has been shown to be a reservoir for several intracellular bacteria pathogens [8, 13, 16, 17].

*Acanthamoebae spp*., are widely used in research as non-mammalian surrogate organisms to study host-interactions [20], including *C. jejuni* [11, 12, 21]. Studies have primarily focused on *C. jejuni*-amoebal interaction population measurements, which enumerate surviving bacteria by colony-forming units, but there is little information at the single-cell level. Some studies have reported on the ability of *C. jejuni* to be phagocytosed by amoebae and thrive within this niche, where they multiply and “burst” out, leading to the release of *C. jejuni* into the environment [11]. This observation is seen in many other bacteria pathogens [14]. However, in our previous study we observed uptake of *C. jejuni* by *Acanthamoebae spp*., but not proliferation [12]. Instead, we observed that *Acanthamoebae spp*. are a transient host for *C. jejuni*, and survival of this bacterium within amoebae enhances its subsequent invasion and survivability in human epithelial cells and amoebae cells [12].

Here, we scrutinize the interaction of *C. jejuni* and *amoebae* using single-cell imaging complemented with intra-amoebal *C. jejuni* transcriptome profiling. We investigate the interaction of *C. jejuni* 11168H with *A. castellanii*, a keratitis strain from the T4 genotype, on the single-cell level by live-cell imaging using confocal microscopy. We show that sub populations of *C. jejuni* cells can (i) resist killing by *A. castellanii*; (ii) are released from *A. castellanii* through exocytosis; and (iii) contrary to previous reports [22], *C. jejuni* can maintain this niche upon amoebal encystment. Given the limited genetic repertoire of *C. jejuni* (genome size ~1.6 Mb) we hypothesized that *C. jejuni* would use similar strategies to survive internalization by amoebae as it would with its warm-blooded hosts. Using RNA-Seq we show that *C. jejuni* indeed utilizes similar strategies to survive and persist within amoebae as it does with its warm-blooded hosts. Our data reinforce the observation that *Acanthamoebae spp*. are an important transient host for *C. jejuni* and may enhance environmental persistence of this prevalent pathogen.

## Methods

### Strains and Cultures

Bacteria were stored using protect bacterial preservers (Technical Service Consultants, Heywood, U.K.) at −80 °C. *C. jejuni* strains were streaked on blood agar (BA) plates containing Columbia agar base (Oxoid) supplemented with 7% (v/v) horse blood (TCS Microbiology, UK) and grown at 37 °C in a microaerobic chamber (Don Whitley Scientific, UK), containing 85% N_2_, 10% CO_2_, and 5% O_2_ for 48 hrs. *C. jejuni* strains were grown on CBA plates for a further 16 hrs at 37 °C prior to use.

*Acanthamoeba castellanii* (T4 genotype) Culture Collection of Algae and protozoa (CCAP) 1501/10 (Scottish Marine Institute) were grown to confluence at 28°C in 75-cm^2^ tissue culture flasks containing peptone yeast and glucose (PYG) media (Proteose peptone 20 g, Glucose 18 g, Yeast extract 2 g, Sodium citrate dihydrate 1 g, MgSO_4_ x 7H_2_O 0.98 g, Na_2_HPO_4_ x 7H_2_O 0.355 g, KH_2_PO_4_ 0.34 g in distilled water to make 1000 ml, pH was adjusted to 6.5). Amoebae were harvested by scraping the cells into suspension, and viability was determined by staining with trypan blue and counting by a hemocytometer using inverted light microscopy.

### *C. jejuni* mutant transformation

*C. jejuni* 11168H mutant constructs were obtained from the Campylobacter resource facility (http://crf.lshtm.ac.uk/wren_mutants.htm) and strain 11168H was naturally transformed using the previously described biphasic method [23] with some minor changes. Briefly, *C. jejuni* was cultured on Mueller Hinton (MH) agar for 16 hrs, cells were resuspended in MH broth to an OD_600nm_ = 0.5, 0.5 ml of this suspension was added to a 15 ml falcon that contained 1 ml of MH agar, *C. jejuni* donor DNA 0.1 – 1 μg was added to the suspension and incubated for 24 hrs at 37 °C in microaerobic conditions, the suspension was plated on MH agar with respective antibiotic for up to 5 days. *C. jejuni* green fluorescent producing bacteria (GFP) strain 11168H_GFP_ was constructed as previously described [12, 24]. *C. jejuni* constructs 11168HΔ*cgb*, 1168HΔ*dsbA* and 1168HΔ*dsbB* were a gift from Dr. Aidan. J. Taylor (University of Sheffield). All strains used in the study can be found in Supplementary File 1 **(Table S2)**.

### *C. jejuni* invasion and survival assay

Invasion and survival assays were performed as previously described [12, 25], with minor alterations. Briefly, *C. jejuni* 11168H, a derivative of the original sequence strain NCTC 11168, and its mutants was incubated with a monolayer of approximately 10^6^ *A. castellanii* at a multiplicity of infection (M.O.I) of 200:1 for 3 hrs at 25 °C in 2 ml PYG media (the M.O.I. of 200:1 was chosen to allow maximal internalization of bacteria without impacting amoebae predation through density-dependent inhibition). The monolayer was washed 3x with phosphate buffer saline solution (PBS) and incubated for 1 hr in 2 ml of PYG media containing 100 μg/mL of gentamicin. *C. jejuni* cells were harvested by scraping the amoebae into suspension and centrifuged for 10 mins at 350 x g to pellet the bacteria and amoebae. Supernatant was discarded and the pellet was suspended in 2 ml of distilled water containing 0.1% (v/v) Triton X-100 for 10 mins at room temperature with vigorous pipetting (every 2 mins) to lyse the amoebae and release bacteria cells. The suspension was then centrifuged for a further 10 mins at 4000 x g, the resultant pellet was resuspended in 1 mL PBS and enumerated for colony forming units on CBA plates for up to 72 hrs at 37 °C microaerobically.

### Live-cell imaging

Green fluorescent protein (GFP) producing *C. Jejuni* 11168H_GFP_ strain was used for confocal live-cell imaging. Time lapse experiments were performed as described above with minor changes. *A. castellanii* cells were incubated in PYG media supplemented with Dextran conjugated-Texas Red (10000 MW; Thermofischer) at a final concentration of 100 μg/ml overnight. *A. castellanii* cells were washed three times and approximately 10^6^ of adherent amoebae were infected with *C. jejuni* 11168H_GFP_ (M.O.I of 200) in 35 mm *μ*-Dish devices (IBIDI) prior to imaging. Texas Red-conjugated dextran was monitored in the red region (excitation and emission wavelengths are 595/615 nm) and GFP was monitored in the green region (excitation and emission wavelengths are 488/510 nm).

Confocal laser scanning microscopy images were obtained using an inverted Zeiss LSM 880 confocal microscope (Zeiss, Germany). Images were taken at 5 sec intervals for time lapse experiments.

### RNA-Seq and real-time RT-qPCR

RNA-Seq was used to identify differentially expressed genes between intra-amoebal *C. jejuni* strain 11168H and the control (which was incubated in the absence of *A. castellanii*) after a total of 4 hrs. Experiments were performed as described above, briefly, *A. castellanii* 10^6^ were infected co-incubated with *C. jejuni* 11168H (M.O.I of 200) for 3 hrs and washed three times to remove extracellular bacteria. Gentamycin was added to a final concentration of 100 μg/ml, this culture was incubated for a further 1 hr before washing three times and RNA was extracted using Triazole (Sigma Aldrich) following manufactures protocol. Ribosomal RNA was depleted using Ribominus (Invitrogen) and libraries was prepared using TruSeq^®^ Stranded mRNA (Illumina). Raw reads were obtained from an Illumina MiSeq paired-end sequencing platform (Illumina). The paired-end reads were trimmed and filtered using Sickle v1.200 [26], Bowtie2 [27] was used to map the reads against the reference sequence; *C. jejuni* strains 11168H assembly GCA_900117385.1. Cufflinks suite [28] was used to convert annotations from GFF to GTF format and Bedtools [29] was used to generate transcript counts per samples. Statistical analysis was performed in R using the combined data generated from the bioinformatics as well as meta data associated with the study (multifactorial design). Adjusted *p*-value significance cut-off of 0.001 and log fold change cut-off of >1.5 was used for multiple comparison.

Expression of genes of interest were quantified by real-time RT-qPCR and normalized against *gyrA*. A 1 μg of total RNA of each sample was reverse-transcribed to cDNA using RT^2^ first strand kit (Qiagen) according to manufactures protocol. Quantification of gene expression was achieved by real-time RT-qPCR using Sybr™ Green real-time PCR master mix using primers generated using PrimerQuest (IDT) (Supplementary File 1 **(Table S3)**). Real-time RT-PCR was performed in 96-well plates using an ABI PRISM 7300 Real-time PCR System (Applied Biosystems) and the relative gene expression for the different genes was calculated from the crossing threshold (Ct) value according to the manufacturer’s protocol (2-ΔΔCt) after normalization using the *gyrA* endogenous control (68).

### Statistical analysis

Evaluation of *C. jejuni* interaction with *A. castellanii* was based on at least 500 amoebae cells per experiment. All experiments presented are at least three independent biological replicates. RNA-Seq analysis were performed in R using the combined data generated from the bioinformatics. Differentially expressed genes were considered significant when the *p*-value of three independent biological experiments was below 0.001. All other data were analyzed using Prism statistical software (Version 9, GraphPad Software) and statistical significance was considered when the *p*-value was <0.05. All values are presented as standard deviation of at least three independent experiments.

### Data availability

The data that supports the RNA-Seq findings of this study are available in the Gene expression Omnibus (GEO) under data set identifier GSE206909.

## Results

### Undigested *C. jejuni* are released back into the environment

As a predator, free-living amoebae inhabits aquatic environments, a niche often occupied by *C. jejuni* [9, 19, 20]. It is plausible that these two organisms have frequent encounters and have co-evolved. Using green fluorescent protein (GFP) -producing *C. jejuni* strain 11168H (1168H_GFP_) and indigestible-conjugated dextran to track phago/-endosomal pathway, time-lapse imaging (0 – 3.5 hrs post infection (h.p.i)), we studied the interaction of *C. jejuni* and *A. castellanii* at single-cell level. We observed active internalization, trafficking of bacteria into digestive vacuoles and the persistence of undigested *C. jejuni* **(Figure 1)**.

**Figure 1.**
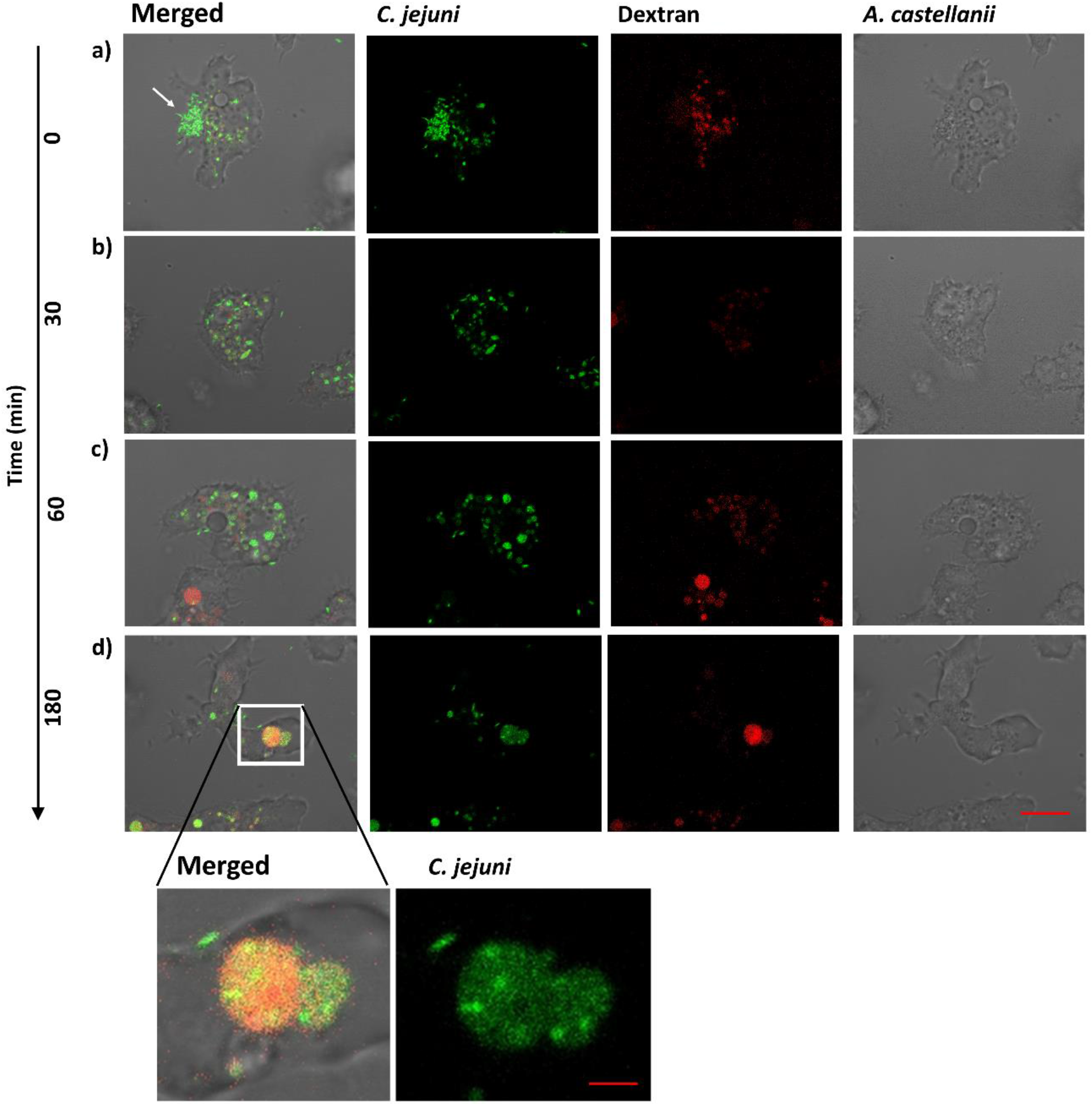
Single cell time-lapse imaging with confocal microscopy monitoring the interaction of *A. castellanii* and *C. jejuni*. **a)** uptake of *C. jejuni* by *A. castellanii* (indicated by arrow); **b)** internalized *C. jejuni* form small bacteria filled vacuoles; **c)** the small bacteria filled vacuoles fuse to form larger vacuoles and **d)** *C. jejuni* colocalizes with Texas Red-conjugated dextran within vacuolar compartments of *A. castellanii* at ~3 h.p.i. (inset; shows fusion of bacterial filled vacuole with dextran filled vacuole). The amoebae are visible in the transmitted-light channel (labeled: *A. castellanii*); GFP-producing *C. jejuni* 11168H_GFP_ in the green channel (labeled: *C. jejuni*); merged signals are shown on the right (labeled: Merged); dextran filled vacuole labeled with Texas Red-conjugated dextran in the red channel (labeled: Dextran). Representation of three independent biological replicates; images were captured with x 63 oil objective, 5 sec intervals. Time is indicated on the left (minutes). Scale bar: 10 *μ*m and 2 *μ*m for inset.

To identify the compartment containing intracellular *C. jejuni*, we followed the time-course of the phago-/endosomal pathway by incubating amoebae for 16 hrs with indigestible fluorophore conjugated to a complex branched glucan, dextran-Texas Red, and tracked localization. Phagocytosed bacteria **(Figure 1a)** formed small bacteria-filled vacuoles **(Figure 1b)** which eventually fused to form larger compartments that colocalized with dextran, this indicated that intracellular *C. jejuni* are trafficked into “digestive-vacuoles” **(Figure 1c and d)**.

Whilst most of the *C. jejuni* were lysed within these compartments, a fraction of the bacteria survived. This was based on retained integrity of the classical helical-shape morphology of *C. jejuni*. Next, we sought to observe the fate of these intact *C. jejuni* cells within these compartments **(Figure 2)**.

**Figure 2.**
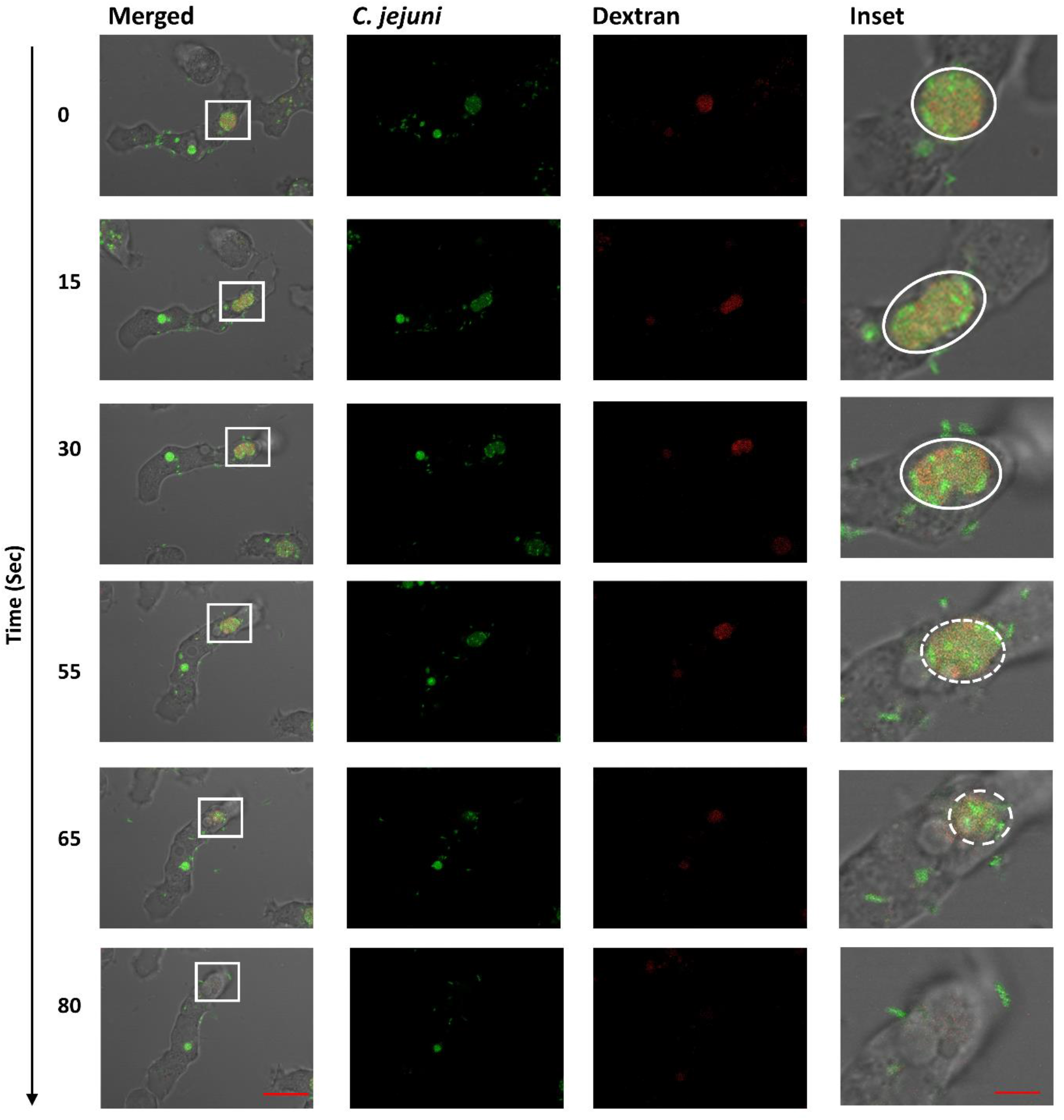
Time-lapse imaging with confocal microscopy showing the release of *C. jejuni* colocalized with dextran. GFP-producing bacteria were incubated with *A. castellanii* for 3.5 hrs and then monitored over a period of 80 sec (relative time shown on the left). Images represent merged transmitted-light (amoebae); GFP channels (*C. jejuni*); red channels (dextran). The inset shows magnification of the white-boxed, areas highlighted with a circle/oval shows an intact compartment; dashed circle/oval indicate release of vacuole content. Representation of three independent biological replicates; images were captured at x 63 oil objective at 5 sec intervals. Time is indicated on the left (in seconds). Scale bars: 10 μm; and 2 μm for inset image. See **Supplementary Movie S1**.

*A. castellanii* were incubated with *C. jejuni* 11168H_GFP_ for 3.5 h and monitored over a period of 80 sec. Live-cell imaging revealed that these undigested bacteria are released back into the environment **(Figure 2 and Supplementary Movie_S1)**. Similar to other phagocytes [30, 31], *A. castellanii* accumulated cellular waste such digested bacteria and the dextran are excreted out of the cell, thus also releasing undigested bacteria back into the environment. We did not observe this phenotype using a non-pathogenic *E*. coli strain DH5α tagged with the fluorophore mCherry (**Supplementary file, Figure S2**).

### *C. jejuni* maintains a niche within amoebic cyst

Contrary to previous reports [11, 21, 25], we did not observe intracellular multiplication of *C. jejuni* and lysis of amoebae, but we did observe *C. jejuni 11168H_GFP_* within the cyst form of *A. castellanii* **(Figure 3)**. *C. jejuni 11168H_GFP_* was incubated with *A. castellanii* for 72 hrs at 25°C in PYG media before imaging with confocal microscopy. Out of 500 amoebic cysts that were enumerated in each biological replicate, an average of ~30% contained GFP-producing bacteria **(Figure 3a and b)**. However, we were unable to recover viable colony forming units (cfu) from this culture.

**Figure 3.**
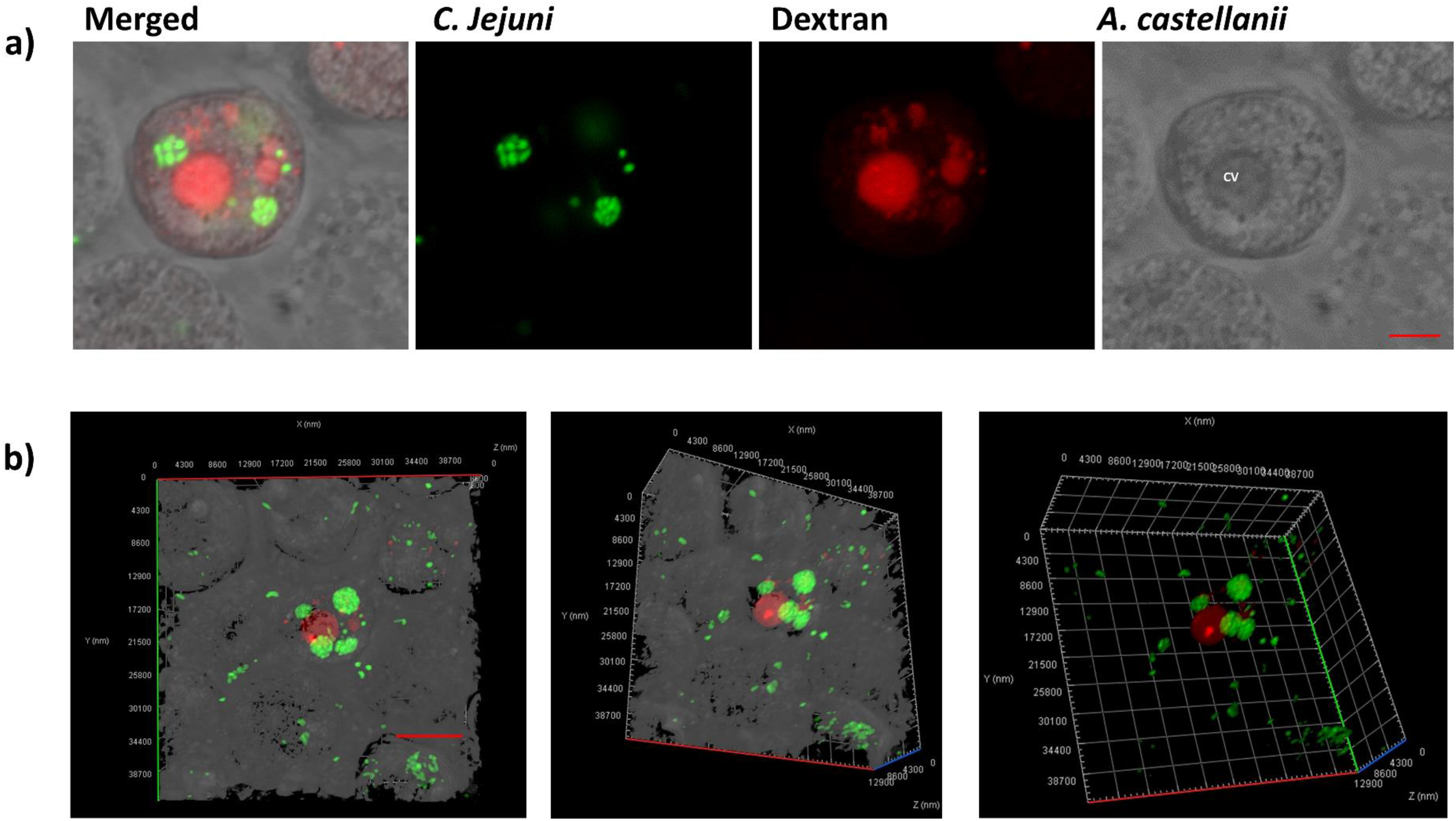
*C. jejuni 11168H_GFP_* within *A. castellanii* cyst. GFP-producing bacteria were incubated with *A. castellanii* (M.O.I 1:200) for 72 hrs and imaged with confocal microscopy, **a)** internalized *C. jejuni* are localized in small tightly packed compartments. The amoebae are visible in the transmitted-light channel (labeled: *A. castellanii*); GFP-producing *C. jejuni* 11168H_GFP_ in the green channel (labeled: *C. jejuni*); merged signals (labeled: Merged); and Texas Red-conjugated dextran filled compartments are shown with the Red channel (labeled: Dextran); CV = contractile vacuole; **b)** volume view of confocal Z-stack showing tightly packed compartments filled with GFP-producing *C. jejuni* (Green); and Texas Red-conjugated dextran filled compartment. Images were captured with x63 oil objective Scale bar: 5 *μm*. A representation of three biological replicate, n=500 amoebic cysts were counted per experiment.

### Intra-amoebal *C. jejuni* transcriptome

To identify the key regulatory changes triggered by survival of *C. jejuni* within *A. castellanii*, RNA-Seq was used to determine differentially expressed genes in strain 11168h intra-amoebae relative to the control at 4 hrs post infection. There was a large difference in gene expression between intra-amoebae *C. jejuni* and the control; a total of 94 genes were differentially transcribed (>1.5-fold; *p*-value <0.001), 72 were up-regulated whilst 22 were down-regulated in the intra-amoebae *C. jejuni* **(Table 1)**.

**Table 1:**
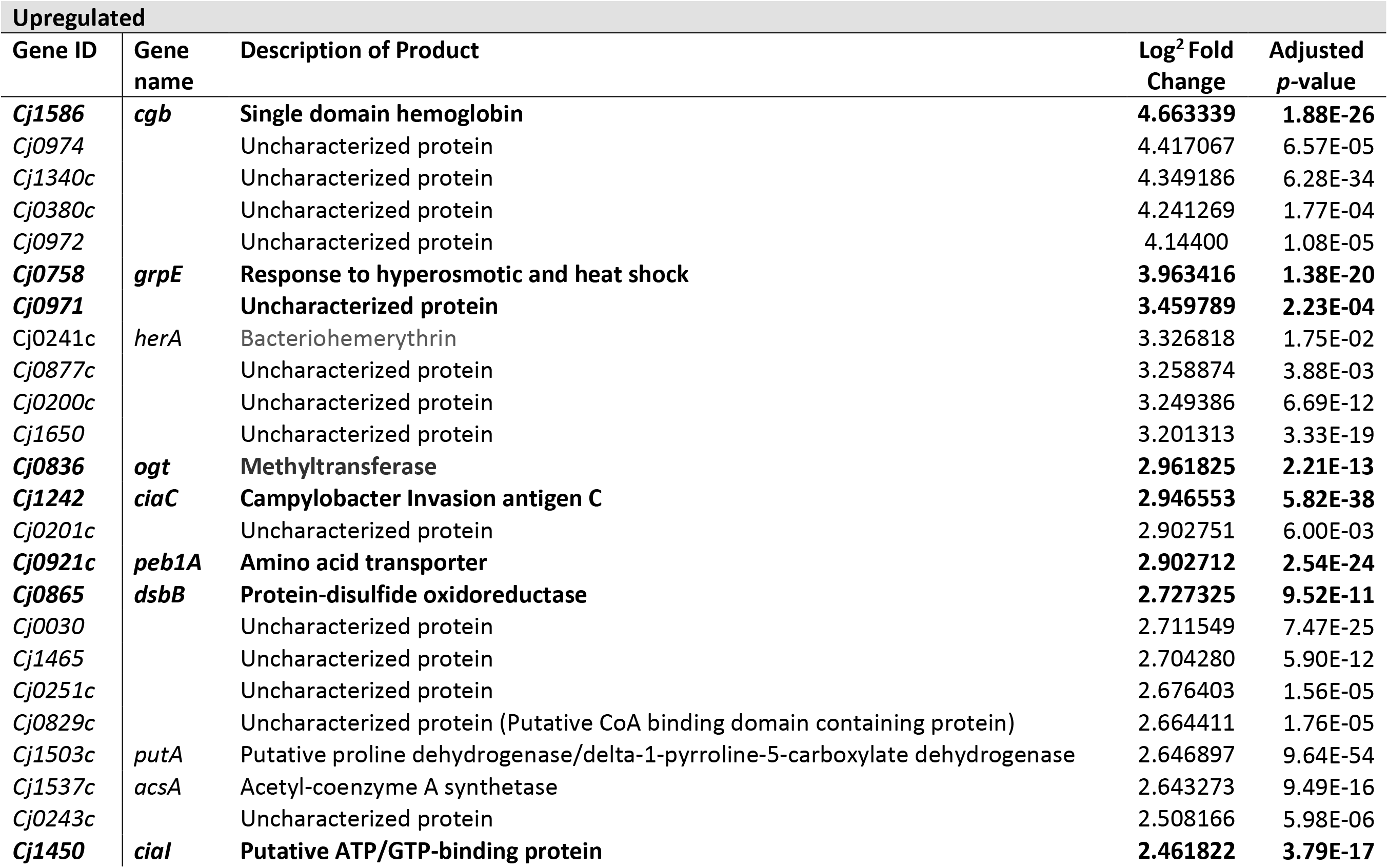
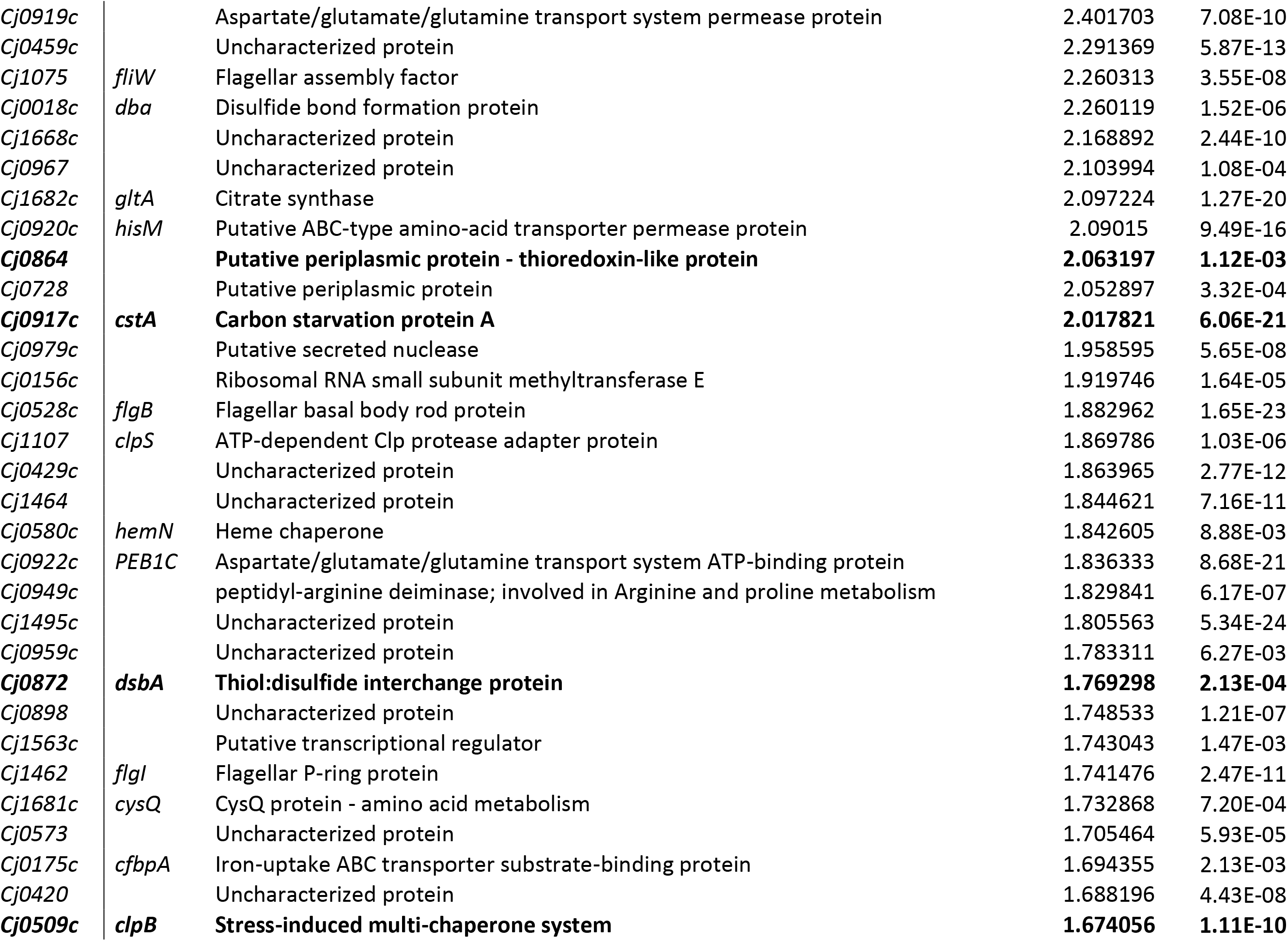
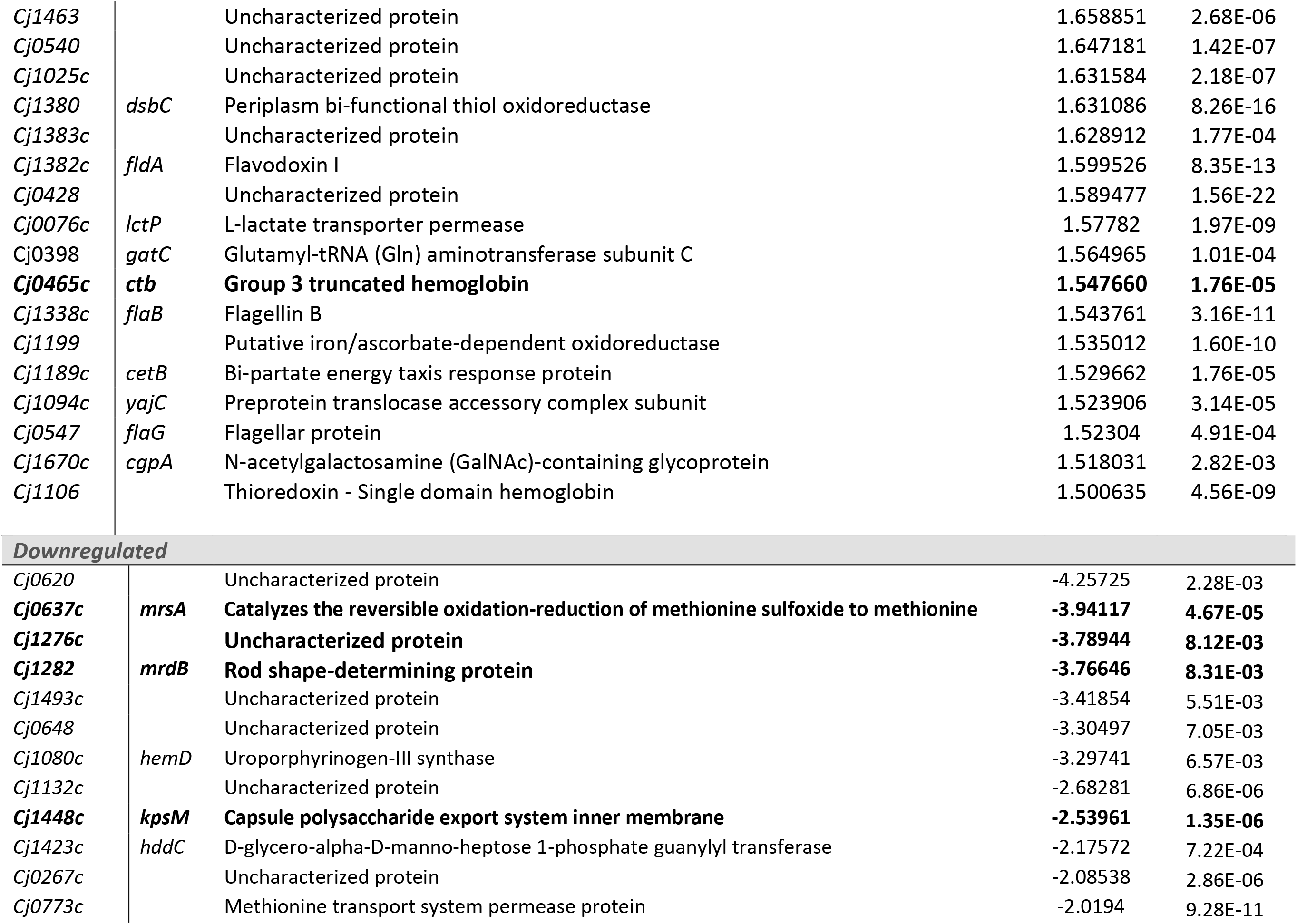
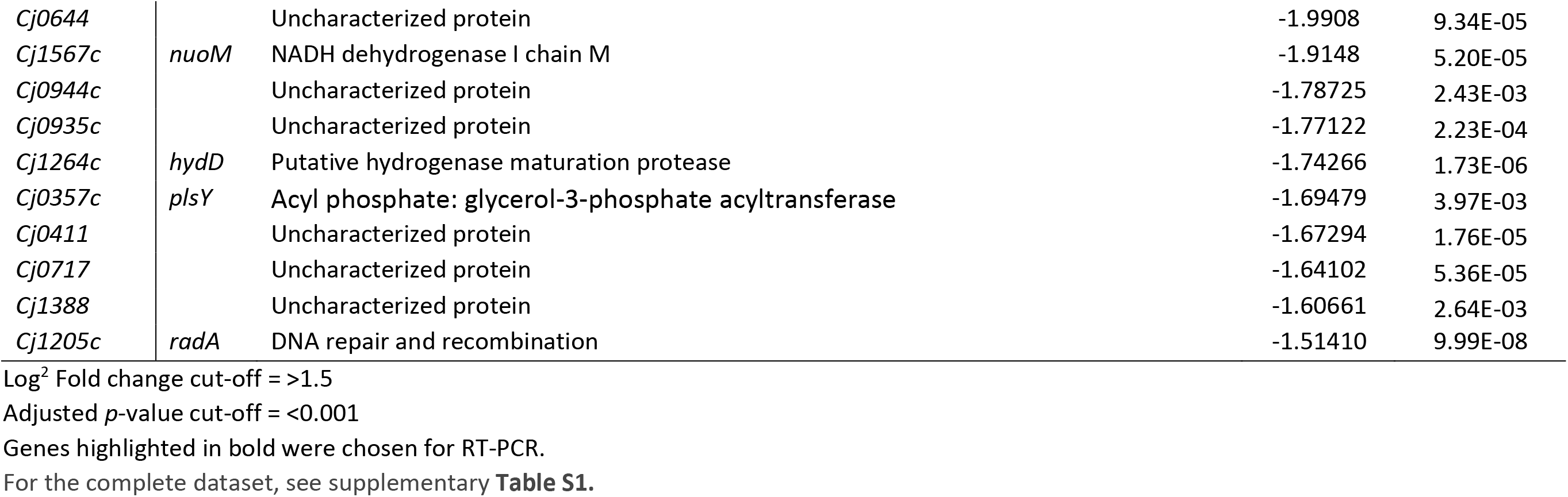
Significantly differentially expressed genes in intra-amoebae *C. jejuni* compared to the control in strain 11168H after 4 hrs post infection.

To verify our RNA-Seq results, RNA was extracted independently of the RNA-Seq experiments for real-time RT-PCR. The first genes were *cgb* which encode a single-domain hemoglobin and *ctb* which encode a truncated hemoglobin from group III of globins; both have a major role in mediating protection of *C. jejuni* against nitric oxide and nitrosative stress [32]. *dsbA* and *dsbB*, both of which encode periplasmic thiol-oxidoreductases that repair disulfide bond in proteins that have been damaged under stress [33–35]; *cstA*, encode carbon starvation protein A; *clpB*, encode an ATP-dependent protease and *grpE*, encode a heat-shock protein, all of which are involved in *C. jejuni* stress response and niche adaptation [36, 37]. We also determined the expression of *peb1A*, which encodes an amino-acid binding protein in addition to facilitating cell adhesion [38], and the expression of *Cj0971*, which encodes an uncharacterized protein. A BLAST (https://blast.ncbi.nlm.nih.gov/) search of *Cj0971* protein sequence revealed the product of *Cj0971* as a filamentous hemagglutinin, this product mediates adhesion and invasion into host cells in other bacteria [39].

We also determined the expression of *ciaC* and *ciaI*, encoding Campylobacter invasion antigens, both of which are essential for invasion and survival within host cells *in vitro* [40, 41]. *KpsM*, encodes transport permease protein and its mutation leads to non-encapsulated C. *jejuni* [42]; *mrsA*, encodes a methionine sulfoxide reductase, involved in oxidative stress repair of proteins containing methionine [43]; *mrdB*, encodes a rod determining protein (RodA). Mutation to *rodA* leads to loss of rod shape in bacilli bacteria [44]; and *Cj1276c*, which encodes an uncharacterized protein. *cgb* and *ctb* had significantly increased transcripts in the RNA-Seq results, real-time RT-PCR indicated a significant (*p*<0.05) ~4.12-fold and a non-statistically significant ~1.95-fold increase in expression, respectively. The two thiol-oxidoreductases (*dsbA* and *dsbB*) also showed significant increase in expression in the RT-PCR experiments (~2.74-fold and ~2.25-fold, respectively) in line with the RNA-Seq results. *cstA, clpB* and *grpE* were all significantly transcribed in the RNA-Seq results, and RT-PCR showed a significant increased expression of *cstA* (~2.74-fold), *grpE* (~2.39-fold) and a non-statistically significant increase in expression of *clpB* (~1.94-fold). In line with our RNA-Seq results, *peb1A* (~2.79-fold), *ciaC* (~3.14-fold), *ciaI* (~2.10-fold) and *Cj0971*(~2.29-fold) were all significantly expressed. *kpsM* (~−2.58-fold), *mrsA* (~−1.41-fold), *mrdB* (~−2.55-fold) and *Cj1276c* (~2.44-fold) were all significantly downregulated in our RT-PCR results, this was reflective of our RNA-Seq results **(Figure 4a)**.

**Figure 4a.**
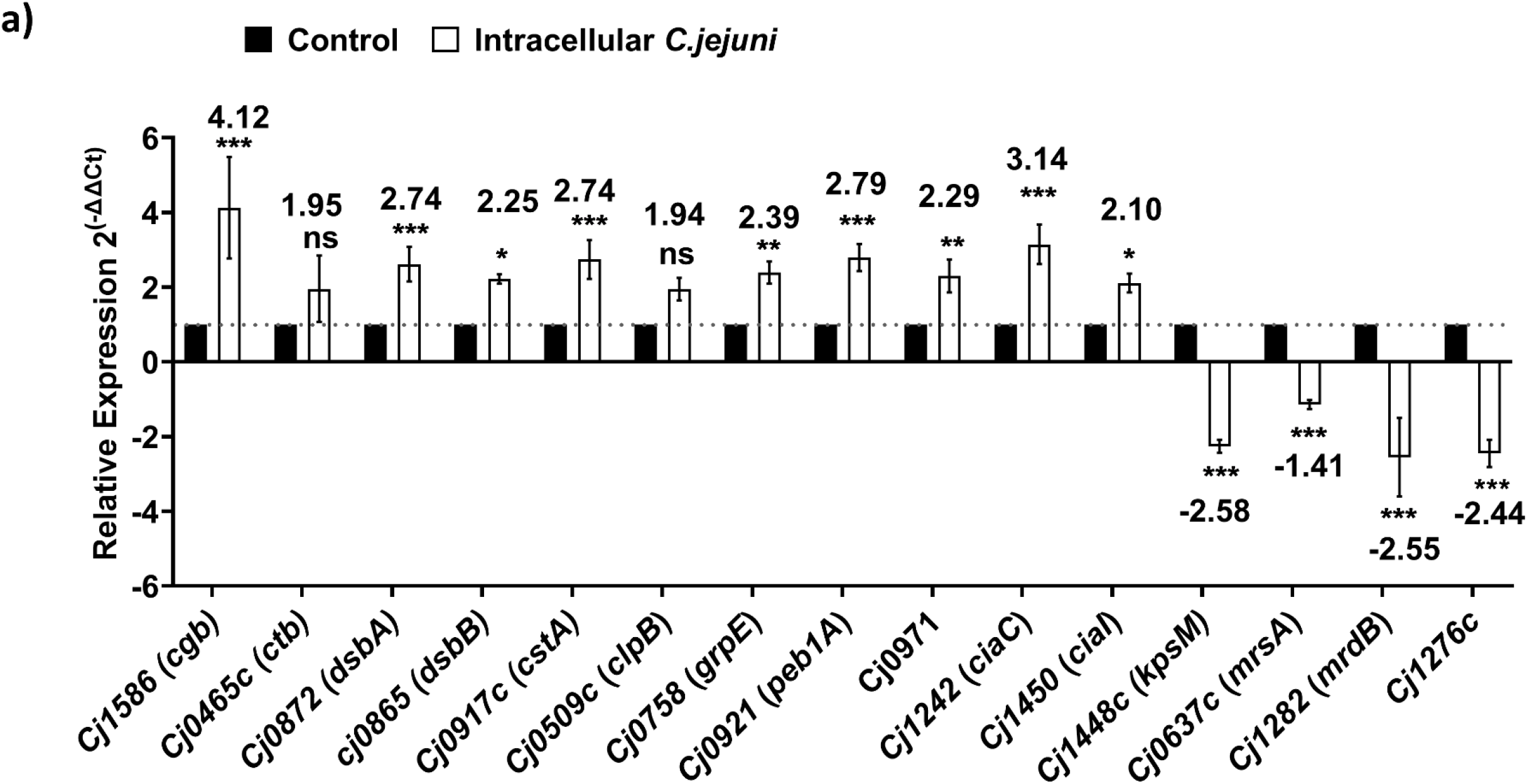
Relative gene expression of *cgb; ctb; dsbA; dsbB; cstA; clpB; grpE; peb1A*; cj0971; *ciaC; ciaI; kpsM; mrsA; mrdB* and *cj1276c*. Expression was determined by real-time RT-PCR and is displayed relative to the control, after normalization using expression of *gyrA*. The values are the means of three independent experiments. Error bars indicate standard deviation., \**p* ≤ 0.05, \*\**p* ≤ 0.01, \*\*\**p* ≤ 0.001. For primers used for RT-PCR see supplementary **Table S3**

Our RNA-Seq data revealed that *C. jejuni* utilizes similar mechanisms to interact with both *A. castellanii* and warm-blooded host cells. Therefore, we generated 10 mutants in *C. jejuni* strain 11168H (**Supplementary File 1: Table S2**) that showed differential expression from our newly generated RNA-Seq data and were previously reported to have a role in interaction of *C. jejuni* with its natural warm-blooded hosts. We then tested their roles in survival within *A. castellanii* by enumerating CFU at 4 hrs post infection **(Figure 4b)**. We chose to test *cgb, clpB, cstA, flgB, dsbA*, *dsbB*, *dsbC*, *flaA* (although this gene did not make the >1.5-fold cutoff, it was upregulated in our RNA-seq by ~ 1.46-fold), *flaB* and *kpsM* mutants.

**Figure 4b:**
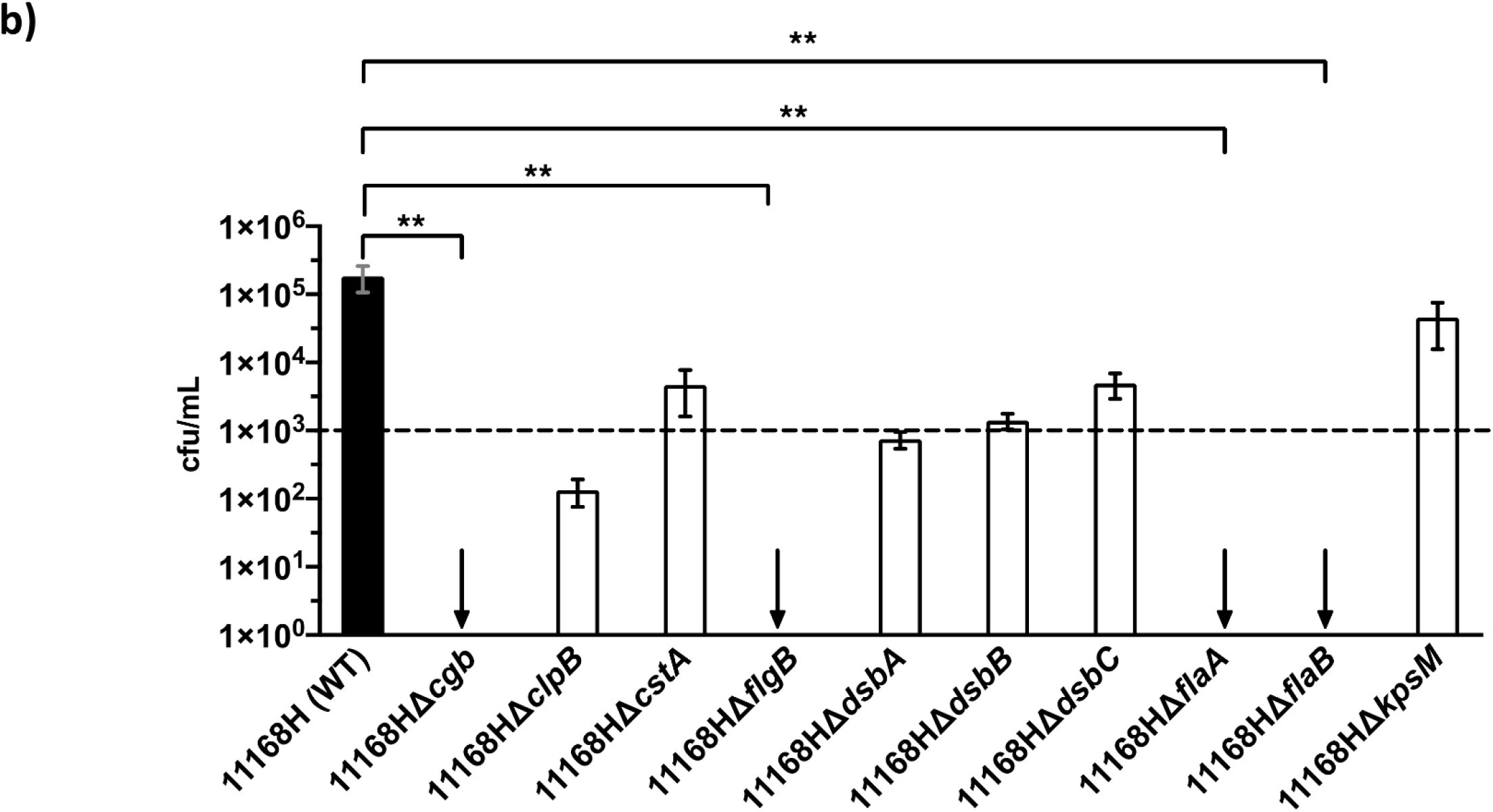
Survival of *Campylobacter jejuni* 11168H and respective mutants within *Acanthamoeba castellanii*. Amoebae were incubated with bacteria at an M.O.I of 1:200 for 3 hrs before treatment with 100 μg/mL of gentamycin for 1 hr. Amoebae were lysed for enumeration of live bacteria. Data is presented as cfu/mL after 4 hrs intracellular survival; black bar = 11168H WT; and white bar = mutants. Error bars represent SD from three independent experiments. One-way ANOVA multiple comparison was used to test for significance; *p < 0.001. dotted line indicates detection limit. Complete data set which includes cfu of inoculum is presented in the supplementary file 1 **(Figure S1a)**.

All mutants tested showed reduced intracellular survivability within *A. castellanii* relative to the 11168H parent strain. Interestingly, mutation to *cgb, flgB* (Flagellar basal body rod protein), *flaA* (major flagellin) and *flaB* (minor flagellin) showed no survival within *A. castellanii*. To test whether this reduction was influenced by amoebal uptake, we enumerated bacteria prior to gentamycin treatment (**supplementary file 1; Figure S1b**). Our results show that whilst 11168HΔ*cgb* uptake by amoebae was similar to the WT levels, mutation to *flgB, flaA* and *flaB* showed reduced uptake by *A. castellanii*, revealing the importance of an intact flagellar during amoebae interaction.

## Discussion

*Campylobacter jejuni* is the leading cause of foodborne gastroenteritis [1, 2]. Understanding its interactions with ubiquitous free-living amoeba in the environment is important and this may shed light on pre-adaptation to survival and virulence mechanisms required when C. jejuni encounters warm-blooded avian or mammalian hosts.

In this study, we investigated the interactions of *C. jejuni* with *A. castellanii* at the single-cell level. We present evidence of previously unreported interactions, and propose part of the *C. jejuni’s* evolution and life-cycle is cohabitation with free-living protist **(Figure 5)**. *C. jejuni* enters the trophozoites phago-/endosomal pathway through phagocytosis, a fraction of the bacteria resists amoebal digestion and they are exocytosed. Previously, we reported that *C. jejuni* are more invasive of mammalian cells and more resistant to killing subsequent to transient internalization by *Acanthamoebae spp.*. Our observation reveals that these more persistent bacteria are also capable of withstanding amoebal encystation.

**Figure 5.**
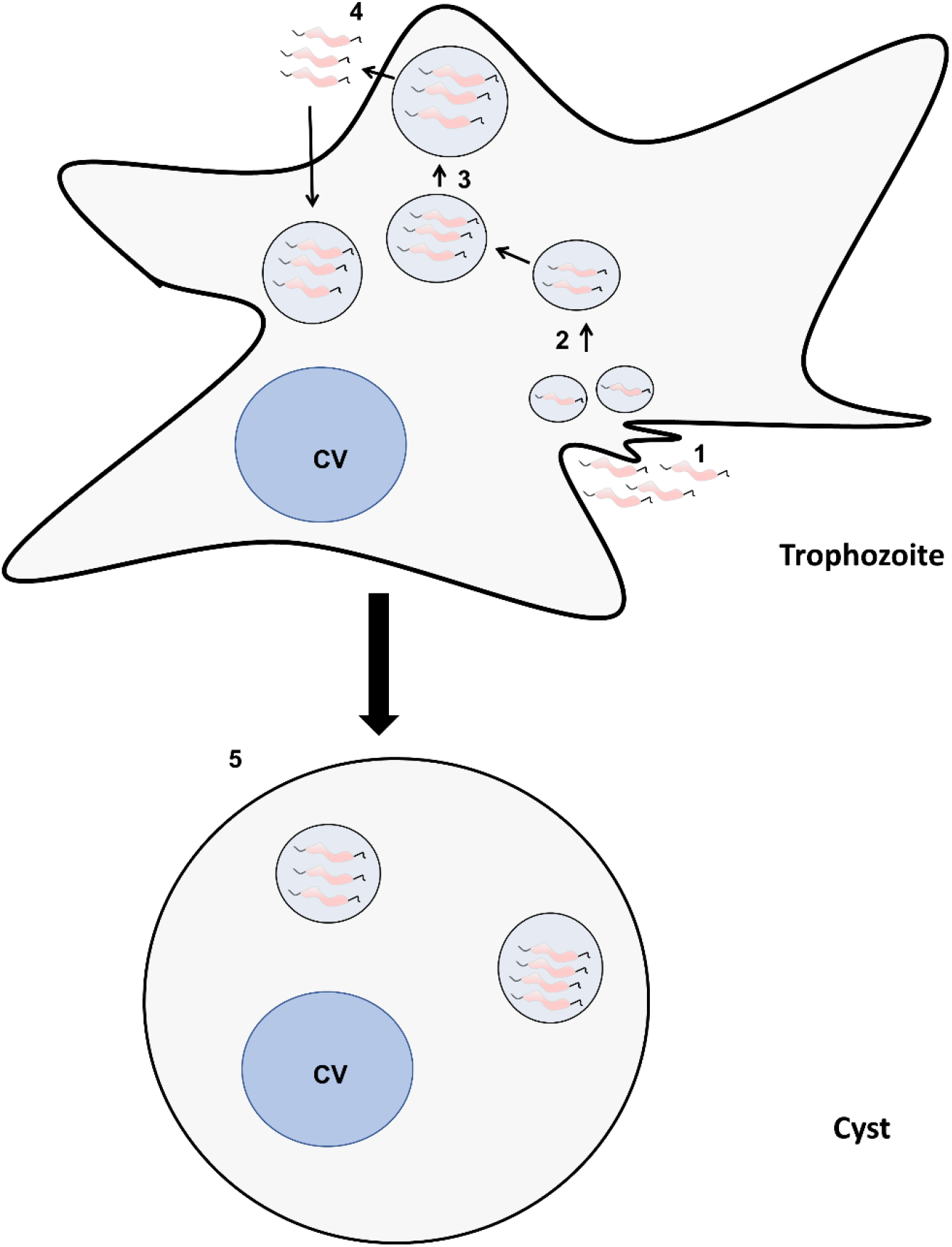
Scheme summarizing the interaction of *C. jejuni* with *A. castellanii*. 1) After phagocytosis by amoebae trophozoites, *C. jejuni* are trafficked into tightly packed vacuoles; 2) which fuse into; 3) larger vacuoles which are emptied through exocytosis; 4) undigested *C. jejuni*, which are more invasive and more resistant to intracellular stresses [12], are capable of re-invading *A. castellanii*; 5) and occasionally these more invasive and resistant *C. jejuni* can withstand encystment of amoebae. CV = contractile vacuole.

*Legionella pneumophila* and *Vibrio cholerae* have long been shown to capably multiply within *Acanthamoebae spp*. and even survive long-term within amoebic cysts [45, 46]. Additionally, *V. cholerae* was shown to multiply within the cysts, which ultimately leads to the destruction of the amoebic cell [46]. Some studies have reported multiplication of *C. jejuni* within amoebae [11, 47] and destruction of amoebic cells through lysis [8]. We did not observe multiplication of *C. jejuni* in our model, however, we observed that long-term co-incubation of *C. jejuni* with amoebae led to transformation of amoebal trophozoites to cysts, which contained GFP producing *C. jejuni*. To our knowledge, this is the first study that shows evidence of *C. jejuni* within *A. castellanii* cysts. The cystic stage of amoebae is known to remain dormant for years and withstand diverse environmental stresses [19, 48]. It is plausible that amoebic cysts may offer *C. jejuni* extra protection from environmental elements, this particular interaction has been reported in *L. pneumophila* [49]. We speculate that following favorable conditions such as transmission to a host (avian or mammalian), this “Trojan horse” would transform into trophozoites potentially leading to the release of its bacteria cargo. However, at this point it is unclear whether encystment is actively induced by *C. jejuni*.

The transient internalization of *C. jejuni* in amoebae has the potential to extend our understanding of how this important pathogen interacts with its natural avian and mammalian hosts. Our transcriptome data of intra-amoebal bacteria revealed that *C. jejuni* utilizes similar factors to interact and survive within amoebae as it does with its natural hosts, adding weight to our previous hypothesis [12]. Similar to the interactions with its natural host [50, 51], the nitrosative stress response genes, *cgb* and *ctb*, were among the most abundant transcripts in intra-amoebal *C. jejuni*, indicating nitric oxide (NO^•^) response. NO^•^ is an antimicrobial agent and an important component of host immune system [52, 53]. Disruption of *C. jejuni cgb* showed significant (*p*<0.05) reduction in survival in our model, emphasizing the importance of NO^•^ defense, but also similarities between *A. castellanii* and *C. jejuni* warm-blooded hosts.

Survival and persistent within a host depends upon sensing stress and responding accordingly [54]. Our data also showed upregulation of stress response in intra-amoebal *C. jejuni*, particularly gene products associated with DnaK, which are involved in suppressing protein aggregation consequently to environmental stress and are important to *C. jejuni* niche adaptation [50, 55–57]. Other upregulated proteostasis genes were of the thiol:disulfide interchange system that repair protein disulfide bonds, including redox stress [58]. This system is also important during *C. jejuni* interaction with its hosts [50, 55, 59, 60]; and mutation to *dsbABC* showed reduced survivability in our amoebae model. Motility and taxis are also indispensable determinants of invasion and survival within *C. jejuni* warm-blooded host [61–63], our RNA-Seq data showed that some of the key genes involved in biosynthesis and maintenance of the flagellar were up-regulated. Disruption of *flgB* and the two heavily O-glycosylated flagellins, *flaA* and *flaB*; the former was shown to be involved in host cell invasion [64], showed decreased invasion to *A. castellanii*. Additionally, mutation to *flgB* was previously shown to be important in the secretion of campylobacter invasion antigens and other modulatory proteins through the putative flagellar-Type 3 Secretion System (FT3SS) [40, 41, 65]. This interesting find suggests that internalization by *A. castellanii* is partly bacterial dependent, and emphasizes the importance of a fully intact and functional flagellar in *C. jejuni*-host interaction [66]. Whether it is motility per se, and/or the non-motility properties of the *C. jejuni* flagellar that are important for these interactions, this remain to be fully evaluated.

*C. jejuni* has a plethora of nutrient uptake and transport systems [50, 55, 67], which are complemented by the central carbon metabolism and electron transport system. We observed differential expression of these genes, indicating that intra-amoebic *C. jejuni* finely tunes its metabolic requirements to adapt to this niche. Both transient and obligate intracellular pathogens are characterized by their differential expression of their metabolic needs [68, 69]. We observed increased up-regulation of metabolic genes in the intra-amoebic *C. jejuni* relative to the control, indicative that the intracellular environment of *A. castellanii* is nutrient-restrictive. This was further supported by the up-regulation of energy taxis signaling response system, *cetB* [55, 70].

We also observed significant differential expression of uncharacterized genes; given that some of these genes were highly upregulated, they might be of functionally importance to the transient intracellular lifestyle of *C. jejuni* and characterization of their products might be key to advancing our understanding of this enigmatic pathogen. The novel observation that amoebae are capable of harboring *C. jejuni* necessitates further investigation on the survival of this pathogen outside the warm-blooded host. This study highlights that amoebae and *C. jejuni* warm-blooded hosts have similar properties which makes *A. castellanii* a useful model to study *C. jejuni-host* interactions.

## Author Contributions

FN conceived the study. FN designed the study. MH performed survival assay; FN performed all other experiments. BL performed RNA-Seq data analysis. FN and BW drafted the manuscript. All authors contributed to editing the manuscript and gave final approval for publication.

## Acknowledgments

We thank Elizabeth McCarthy of the [Wolfson Cell Biology Facility (WCBF)](https://www.lshtm.ac.uk/research/faculties/itd/wolfson-cell-biology-facility) at LSHTM for providing training and technical assistance with the inverted Zeiss LSM880 confocal microscope. We thank Dr. Aidan J Taylor for providing us with mutant constructs and Ana T. López-Jiménez for her advice.

