## Supplementary File 1 for "Survival of *Campylobacter jejuni* in *Acanthamoebae castellanii* provides mechanistic insight into host pathogen interactions"

**Running title:** *Campylobacter jejuni* interactions with *Acanthamoebae castellanii*

Fauzy Nasher[1*]., Burhan Lehri[1]., Megan F Horney[1]., Richard A Stabler[1]., Brendan W Wren[1*].

[1]Faculty of Infectious and Tropical Diseases, London School of Hygiene and Tropical Medicine, London, United Kingdom.

### Figure S1:

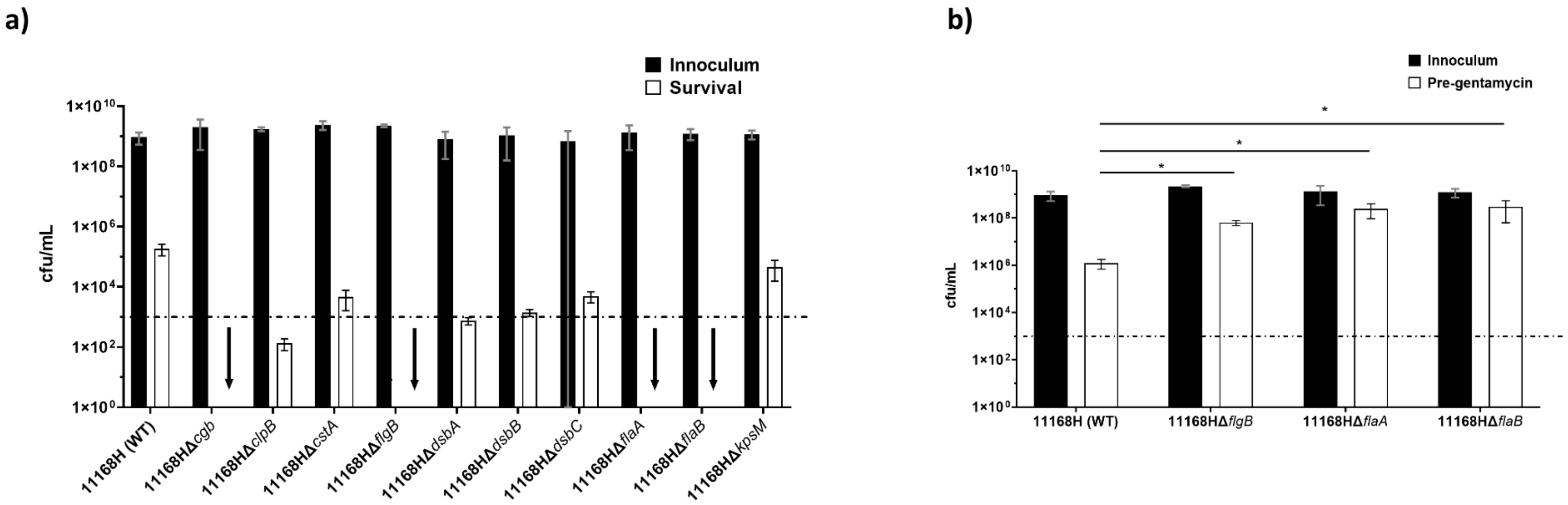


**Figure S1: Survival of *Campylobacter jejuni* 11168H and respective mutants within *Acanthamoeba castellanii*. a)** Amoebae were incubated with bacteria at an M.O.I of ~200:1 for 3 hrs before treatment with 100 µg/mL of gentamycin for 1 hr. Amoebae were lysed for enumeration of live bacteria. b) pre-gentamycin treatment enumeration to check for uptake of *11168H*Δcgb; *11168H*ΔflgB; *11168H*ΔflaA; and *11168H*ΔflaB by *A. castellanii*. Data is presented as cfu/mL; error bars represent SD from three independent experiments. Two-way ANOVA multiple comparison was used to test for significance; $*p \leq 0.05$.

**Figure S2:**

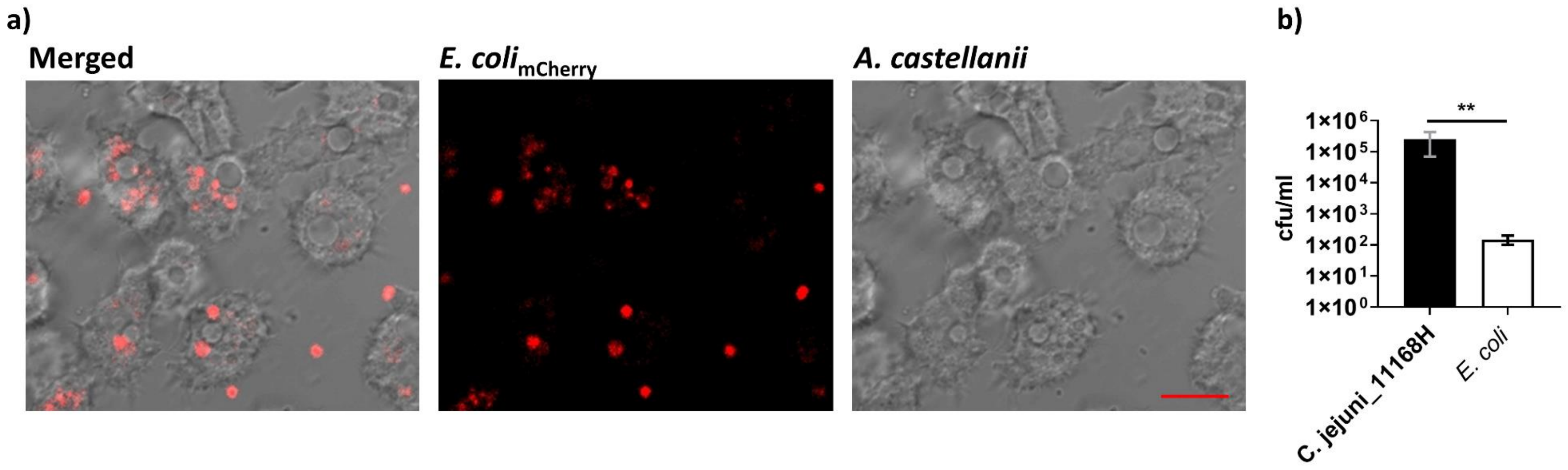


**Figure S2: Survival of *E.*coli Dh5α$_{\mathbf{mCherry}}$ within *A. castellanii*. a)** Confocal microscopy images showing *E. coli*$_{\text{mcherry}}$ within *A. castellanii*; **b)** CFU of *E. coli*$_{\text{mCherry}}$ survival within *A. castellanii* after 4 hr infection compared to *C. jejuni* 11168H. Images represent merged transmitted light (amoebae); red channels (*E. coli*). Images were captured at x63 oil objective; scale bars: 10 µm; mCherry fluorescent was monitored in the red region (excitation and emission wavelengths are 587/610 nm). Error bars represent SD from three independent experiments; student *t*-test was used to test for significance **$p \leq 0.001$. Standard laboratory *E. coli* DH5α was transformed with a commercially available plasmid construct pFPV-mcherry, *E. coli* cells were acquired from New England Biolabs (USA) and transformation was performed according to manufactures protocol and selected with ampicillin.

**Table S2: Table of all the strain used in this study**

| Strain | Antibiotic selection |
|---|---|
| 11168HΔ*cgb* | Kanamycin |
| 11168HΔ*clpB* | Kanamycin |
| 11168HΔ*cstA* | Kanamycin |
| 11168HΔ*dsbA* | Kanamycin |
| 11168HΔ*dsbB* | Kanamycin |
| 11168HΔ*dsbC* | Kanamycin |
| 11168HΔ*flaA* | Kanamycin |
| 11168HΔ*flaB* | Kanamycin |
| 11168HΔ*flgB* | Kanamycin |
| 11168HΔ*kpsM* | Kanamycin |
| $11168H_{GFP}$ | Chloramphenicol |
| 11168H | N/A |

**Table S3: Primers used for RealTime RT-qPCR**

| Gene name | Primers for real-time RT-qPCR |
|---|---|
| ***cgb*** **(*Cj1586*)** | F: 5'- GTTGCCATAACTCATGTTAATTTAGGAG-3'<br>R: 5'- CCAAGCTTTAAGAGTGGCTTCA-3' |
| ***Ctb*** **(*Cj0465c*)** | F: 5' -CTATCATTTGTGCACGCTGTA-3'<br>R: 5' - AGCACTTAGATCTACCTCCTTT-3' |
| ***ciaC*** **(*Cj1242*)** | F: 5'-GCAGATGAATTTCAAGCCACAT-3'<br>R: 5'- TTCTCTAACAGCACCAAGATCAA-3' |
| ***Peb1A*** **(*Cj0921*)** | F: 5'-GCTATCACCGCATCTACACTAC-3'<br>R: 5'- GTTTCGAAGTAGATGTTGCCAAA-3' |
| ***Cj0971*** | F: 5'-CAGCTGATGAACTAACAAGCATTA-3'<br>R: 5'- CAATACTTCCTTTAAAGCGTTTGC-3' |
| ***dsbA*** **(*Cj0872*)** | F: 5' -CTCTATCCTGTAAGTTTAATGAATGGG-3'<br>R: 5'- CTATCAGAATAACTCGCATCTTTACC-3' |
| **(*cstA*) (*Cj0917*)** | F: 5'-CCCAATAAGGATCAGCAGCTATAA-3'<br>R: 5'- TTATCCGTCCAGGTAGAGTAGG-3' |
| ***clpB*** **(*Cj0509c*)** | F: 5'-CGGTTGCTCTACCATCATCTAA-3'<br>R: 5'- CTTACTGAAGCCGTACGAAGAA-3' |
| ***mrdB*** **(*Cj1282*)** | F: 5'-AGCTATAAGTGTTAGCTCTCCTATT-3'<br>R: 5'- TTTCCGGTTAAACCACCATTTC-3' |
| ***kpsM*** **(*Cj1448*)** | F: 5' -GCAAAGTTCTTAAAGGTTCCACA-3'<br>R: 5' - CCTGTTCATTTGCTTGGAGTTT-3' |
| ***mrsA*** **(*Cj0637c*)** | F: 5'-CTTCTATGTTGCCATCACCATTAC-3'<br>R: 5' - AGCCGTATTTGAACGCCTAA-3' |
| ***Cj1276c*** | F: 5' -CCGTATTTACACACAACAAGCA-3'<br>R: 5' - CCAAGAAAGTTTAAAGGCTGTAGAT-3' |
| ***gyrA*** **(*Cj1027c*)** | F: 5' -AGTAATACGTGGCACATCAAATTTACTTCTAAT-3'<br>R: 5'- GCAGAATTAATGAAAGAAATTGCAAGACTTG-3' |
| ***ciaI*** **(*Cj1450*)** | F: 5' -GAAGGCTCTAAGCTCACACAA-3'<br>R: 5' - TGGCTTTAACTCTCCGACTTTAG-3' |
| ***dsbB*** **(*Cj0865*)** | F: 5'-GCAAGATCTATTGTTGTATGTGCTT-3'<br>R: 5'- CACTATAATTATCAGCGTTAGCAATAGG-3' |

**F= Forward primer**
**R= Reverse primer**
